# A reference single-cell transcriptomic atlas of human skeletal muscle tissue reveals bifurcated muscle stem cell populations

**DOI:** 10.1101/2020.01.21.914713

**Authors:** Andrea J. De Micheli, Jason A. Spector, Olivier Elemento, Benjamin D. Cosgrove

**Affiliations:** Meinig School of Biomedical Engineering, Cornell University, Ithaca, NY, 14853, USA; Division of Plastic Surgery, Weill Cornell Medical College, New York, NY, 10021, USA; Englander Institute for Precision Medicine, Weill Cornell Medicine, New York, NY, 10021, USA

**Author notes:** Corresponding author: Benjamin D. Cosgrove, Meinig School of Biomedical Engineering, Cornell University 159 Weill Hall, 526 Campus Road, Ithaca, NY 14853, USA |.

## Abstract

Single-cell RNA-sequencing (scRNA-seq) facilitates the unbiased reconstruction of multicellular tissue systems in health and disease. Here, we present a curated scRNA-seq dataset of human muscle samples from 10 adult donors with diverse anatomical locations. We integrated ∼22,000 single-cell transcriptomes using Scanorama to account for technical and biological variation and resolved 16 distinct populations of muscle-resident cells using unsupervised clustering of the data compendium. These cell populations included muscle stem/progenitor cells (MuSCs), which bifurcated into discrete “quiescent” and “early-activated” MuSC subpopulations. Differential expression analysis identified transcriptional profiles altered in the activated MuSCs including genes associated with ageing, obesity, diabetes, and impaired muscle regeneration, as well as long non-coding RNAs previously undescribed in human myogenic cells. Further, we modeled ligand-receptor cell-communication interactions and observed enrichment of the TWEAK-FN14 pathway in activated MuSCs, a characteristic signature of muscle wasting diseases. In contrast, the quiescent MuSCs have enhanced expression of the *EGFR* receptor, a recognized human MuSC marker. This work provides a new technical resource to examine human muscle tissue heterogeneity and identify potential targets in MuSC diversity and dysregulation in disease contexts.

## Introduction

Skeletal muscles are essential to daily functions such as locomotion, respiration, and metabolism. Upon damage, resident muscle stem cells (MuSCs) repair the tissue in coordination with supporting non-myogenic cell types such as immune cells, fibroblasts, and endothelial cells (Bentzinger et al., 2013). However, with age and disease, the repair capacity of MuSCs declines, leading to complications such as fibrotic scarring, reduced muscle mass and strength (Blau et al., 2015; Järvinen et al., 2014), fat accumulation and decreased insulin sensitivity (Addison et al., 2014), all of which severely affect mobility and quality of life (Larsson et al., 2018).

Human MuSCs are defined by the expression of the paired box family transcription factor PAX7 and can be isolated using various surface marker proteins including β1-integrin (CD29), NCAM (CD56), EGFR, and CD82 to varying purities (Pisani et al., 2010; Charville et al., 2015; Alexander et al., 2016; Uezumi et al., 2016; Wang et al., 2019). With ageing, human MuSCs exhibit a heterogeneous expression of the senescence marker p16^Ink4a^ and accumulate other cell-intrinsic alterations in myogenic gene expression programs, cell cycle control, and metabolic regulation (Sousa-Victor, et al., 2014; Blau, et al., 2015). However, given their varied molecular and functional states, our understanding of MuSCs in adult human muscle tissue remains incompletely defined. In addition, cellular coordination in the regulation of human muscle homeostasis and regeneration remains poorly understood due to the lack of experimentally tractable models with multiple human muscle cell types. Given these challenges we posited that an unbiased single-cell reference atlas of skeletal muscle could provide a useful framework to explore MuSC variability and communication in adult humans.

Here, we deeply profiled the transcriptome of thousands of individual MuSCs and muscle-resident cells from diverse adult human muscle samples using single-cell RNA-sequencing (scRNA-seq). After integrating these donor datasets to conserve biological information and overcome technical variation, we resolved two subpopulations of MuSCs with distinct gene expression signatures. Using differential gene expression analysis and ligand-receptor interaction modeling, we extend the known repertoire of human MuSC gene expression programs, suggesting new regulatory programs that may be associated with human MuSC activation, as well as features of human muscle aging and disease.

## Results

### Collection and integration of a diverse human scRNA-seq dataset

We used scRNA-seq to collect and annotate a single-cell transcriptomic dataset of diverse adult human muscle samples under homeostatic conditions. The muscle samples were from surgically discarded tissue from n=10 donors (range: 41 to 81 years old) undergoing reconstructive procedures and originating from a wide variety of anatomical sites in otherwise healthy patients (**Fig. 1A**). Each sample was ∼50 mg after removal of extraneous fat and connective tissue. Muscle samples were enzymatically digested into single-cell suspensions and independently loaded into the 10X Chromium system. All together, we collected over 22,000 human muscle single-cell transcriptomes (2206 ± 1961 cells per dataset) into a single data compendium. Using unsupervised clustering, we resolved 16 types of cells of immune, vascular, and stromal origin, as well as two distinct subpopulations of MuSCs and some myofiber myonuclei (**Fig. 1B**).

**Figure 1.**
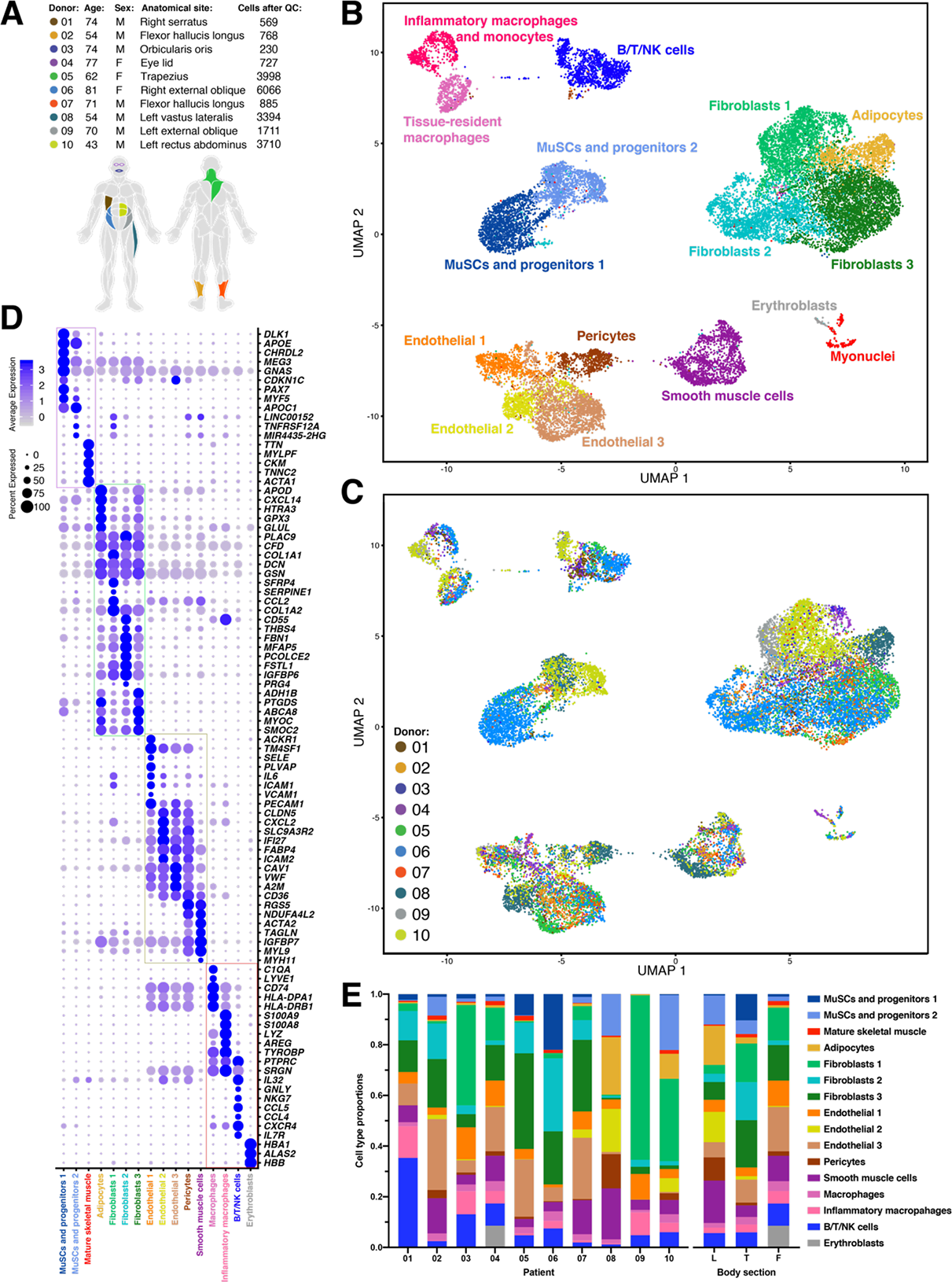
Single-cell transcriptomic map of human muscle tissue biopsies. **(A)** Metadata (sex, age, anatomical site, and the number of single-cell transcriptomes after quality control (QC) filtering) from n=10 donors. Colors indicate sample anatomical sites. **(B)** Scanorama-integrated and batch-effect corrected transcriptomic atlas revealing a consensus description of 16 distinct muscle-tissue cell populations. **(C)** Transcriptomic atlas colored by donor and anatomical location. **(D)** Dot-plot showing differentially expressed genes that distinguish the cell populations. Grouped in four compartments: muscle, endothelial/vascular, stromal, and immune. **(E)** Cell type proportions as annotated in (B) across the 10 donors and grouped by body sections. L = leg (donors 02, 07, 08), T = trunk (donors 01, 05, 06, 09, 10), F = face (donors 03, 04).

Given important differences in anatomical site, donor health history, age, sex, and surgical procedures, the muscle samples were highly heterogeneous in terms of cell-type diversity and underlying gene expression profiles. Comparing the resulting scRNA-seq datasets is therefore a challenge that we addressed using recently developed bioinformatic integration methods (Stuart et al., 2019ab; Hie et al., 2019). Our goal was to assemble a unified dataset of human muscle tissue that faithfully conserved sources of biological variability such as donor, anatomical location, and cell composition heterogeneity, while accounting for technical biases. We tested four different scRNA-seq data integration methods (**Fig. S1**) and found that Scanorama (Hie et al., 2019) followed by scaling the output by regressing against the library chemistry technical variable (“10X chemistry”) and the number of genes detected per single-cell best satisfied this goal. Detailed information on our methodology is provided in **Fig. S1**. After integrating the 10 datasets, we noted remarkable consistency amid cell types across donors (**Fig. 1C, 1E**); owing to the robustness of scRNA-seq technology, the bioinformatic method chosen, and our sample preparation protocol. Differential gene expression analysis between the 16 distinct subpopulations identified an extensive set of unique markers that we grouped into 4 categories (**Fig. 1D**).

### scRNA-seq resolves the cellular diversity of human muscle and novel markers

We annotated and interpretation the consensus cell atlas (**Fig. 1B, D**) into cell type sub-populations as follows. We identify four types of stromal cells starting with adipocytes found to be expressing apolipoprotein D (*APOD*) (Muffat et al., 2010), the brown fat tissue adipokine *CXCL14* (Cereijo et al., 2018), *GPX3*, and *GLUL*. Among the 3 other subpopulations of fibroblast-like cells, Fibroblasts 1 express high levels of collagen 1 (*COL1A1*), *SFRP4*, *SERPINE1*, and *CCL2*; Fibroblasts 2 express fibronectin (*FBN1*), the microfibril-associated glycoprotein *MFAP5*, and *CD55* known to be expressed by synoviocytes (Karpus et al., 2015); and Fibroblast 3 is mainly characterized by *SMOC2*, a marker of tendon fibroblasts (Swanson et al., 2019).

We also identify 5 types of vascular cells, including 3 endothelial subpopulations, and a subpopulation of pericytes and smooth muscle cells (SMCs). Pericytes and SMCs express the canonical markers *RGS5* and *MYH11*. Endothelial 1 express E-selectin (*SELE*), *IL6*, *ICAM1*, *VCAM1*. These genes are upregulated at sites of inflammation to facilitate immune cell recruitment, suggesting this Endothelial 1 cell population may be involved in homeostatic muscle tissue remodeling (Watson et al., 1996; Goncharov et al., 2017). Endothelial 2 cells are distinguished by expressing high levels of claudin-5 (*CLDN5*), *ICAM2*, and the chemokine *CXCL2*. Endothelial 3 express high levels of the platelet-recruiting Von Willebrand Factor (*VWF*) and caveolin-1 (*CAV1*), a protein known to regulate cholesterol metabolism, atherosclerosis progression, as well as MuSC activation (Fernández-Hernando et al., 2010, Volonte et al., 2004).

We also noted two types myeloid immune cells. First, tissue-resident and anti-inflammatory macrophages which express *CD74* and histocompatibility complex *HLA* proteins. Second, activated macrophages and monocytes that express inflammatory markers such as *S100A9* (calgranulin) and *LYZ* (lysozyme). Moreover, *S100A9* transcript abundance levels have been shown to be a feature in ageing and chronic inflammation (Swindell et al., 2013). We also identified a pool of T/B lymphocytes and NK cells characterized by *IL7R* and *NKG7,* respectively, as well as a small subset of *HBA1*^+^ erythroblasts.

Finally, we identified two subpopulations of MuSCs (henceforth called “MuSC1” and “MuSC2”). MuSC1 highly expressed the canonical myogenic transcription factor *PAX7* (Kuang et al., 2006), as well as chordin-like protein 2 (*CHRDL2*) and Delta-like non-canonical Notch ligand 1 (*DLK1*). *CHRDL2* has been shown been previously shown to be expressed in freshly isolated quiescent human MuSCs (Charville et al., 2015), though its function is still to be understood. DLK1 is an inhibitor of adipogenesis whose role in muscle has mainly been recognized in the embryo but remains controversial in adult muscle regeneration (Waddell et al., 2010; Andersen et al., 2013; Zhang et al., 2019). In contrast to MuSC1, MuSC2 expressed lower levels of *PAX7* but maintain expression of *MYF5* (a marker of activated MuSCs) and *APOC1* (**Fig. 2B**). Interestingly, the MuSC2 population also had elevated expression of two long non-coding RNAs (lncRNAs), *LINC00152* and *MIR4435-2HG*. LncRNAs are involved in regulating myogenesis (Hagan et al., 2017). Surprisingly, we detected low expression of the myogenic commitment factors *MYOD1* and *MYOG* (**Fig. 2B**), in contrast to scRNA-seq analyses of adult mouse muscle (Dell’Orso et al., 2019; De Micheli et al., 2019). These observations suggest that the MuSC1 and MuSC2 populations are both comprised largely of muscle stem cells, not committed myogenic progenitors. In addition, we noted that “Myonuclei” population (**Fig. 1B**) was enriched for myosin light chain (*MYLFP*), skeletal alpha-actin (*ATCA1*), and troponin C (*TNNC2*), proteins involved in muscle contraction. This multiple-donor scRNA-seq atlas highlights the cellular diversity of human muscle tissue and revealed two distinct MuSC subpopulations along with specific myogenic expression programs.

**Figure 2.**
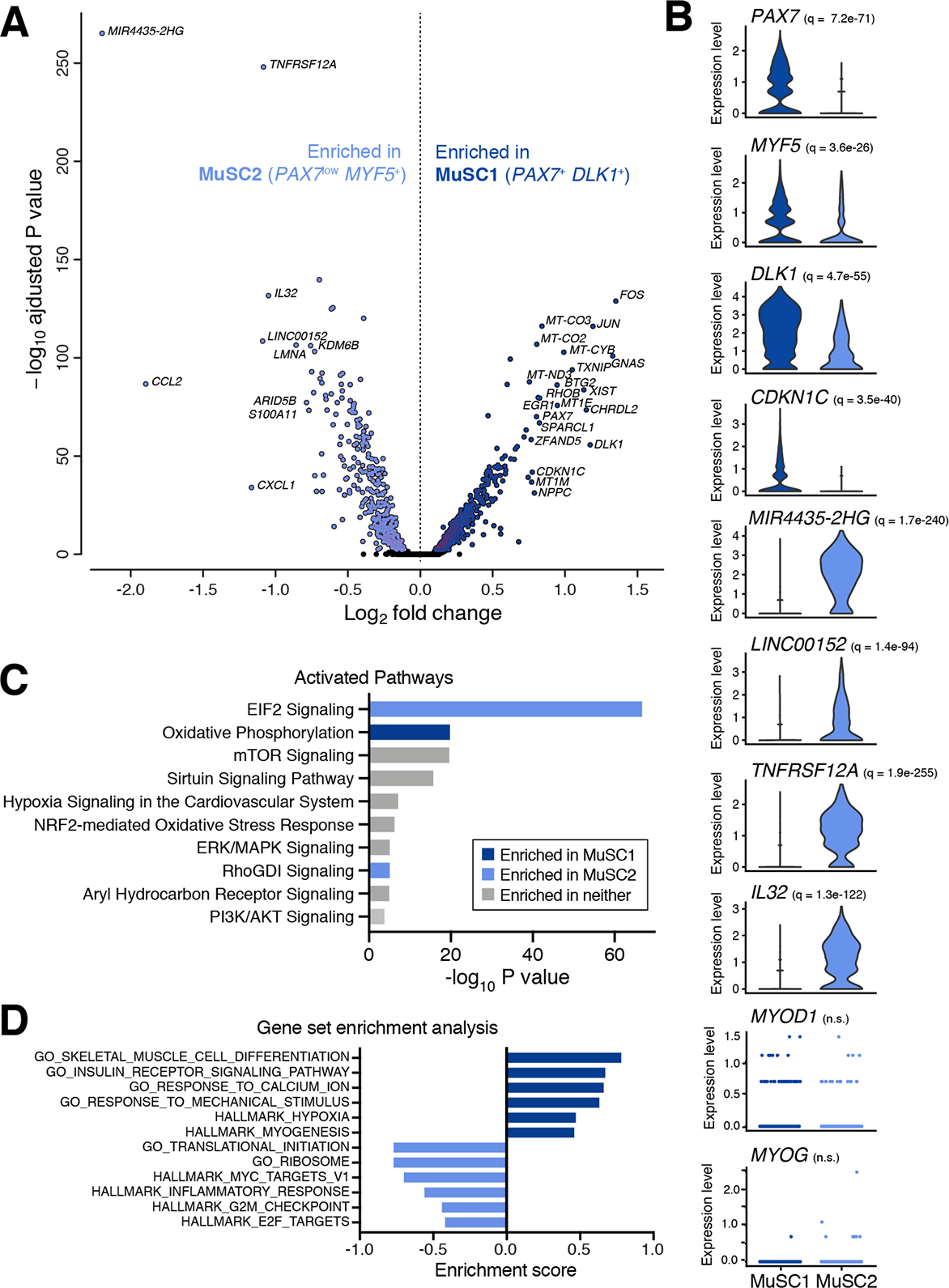
Gene expression and pathway analysis comparison between two MuSC subpopulations. **(A)** Volcano plot from comparing transcript levels between all cells within the “MuSC1” and “MuSC2” subpopulations. Log2 fold-change in normalized gene expression versus -log10-adjusted p-value plotted. Differentially expressed genes (adjusted p value < 0.05) are colored dark or light blue (based on their enrichment in MuSC1 or MuSC2, respectively). Genes with log2 fold-change > 0.75 are labeled. **(B)** Normalized expression values of select differentially expressed genes. *q*-values reported in inset. **(C)** Top activated canonical pathways by Ingenuity Pathway Analysis based on differentially expressed genes and ranked by p value. Pathways significantly enriched in either population with |z-score| > 1 are indicated in blue. **(D)** Select gene ontology (GO) terms and hallmark pathways enriched between the MuSC subpopulations as identified by gene set enrichment analysis (GSEA) and ranked by enrichment score (ES).

### Homeostatic human muscle contains two distinct MuSC subpopulations

Next we examined genes that were differentially expressed between the MuSC1 and MuSC2 subpopulations and the biological processes that characterize them (**Fig. 2A-B**). The MuSC1 subpopulation was enriched for *PAX7*, *DLK1*, and *CHRDL2*, as well as for the cyclin-dependent kinase inhibitor *CKDN1C* (encoding P57^KIP2^), suggesting that these cells are quiescent and not cycling. In addition, this subpopulation expresses the transcription factor *BTG2*, which was identified in mouse to be enriched in quiescent MuSCs (De Micheli et al., 2019). We also note that the MuSC1 subpopulation expressed elevated levels of mitochondrial genes as well as *FOS*, *JUN*, and *ERG1*. Upregulation of these genes has been shown to be potential artefacts of the enzymatic digestion during the sample preparation (van den Brink et al., 2017).

The MuSC2 subpopulation was enriched for multiple markers of inflammation including *CCL2*, *CXCL1*, *IL32*, and surface receptor *TNFRSF12*/*FN14*. In particular, CCL2 and CXCL1 are inflammatory cytokines known to be upregulated in muscle repair, exercise, and fat metabolism (Harmon et al., 1985; Pedersen et al., 2012). In addition, IL32 has been shown to have inflammatory properties in human obesity (Catalán et al., 2017) and have a negative impact on insulin sensitivity and myogenesis (Davegårdh et al., 2017), while TNFRSF12/FN14 has been implicated in various muscle wasting diseases (Mittal et al., 2010; Enwere et al., 2014) and metabolic dysfunction (Sato et al., 2014). Furthermore, the MuSC2 population is enriched for ribosomal gene expression (e.g. *RPLP1* and *RPS6*; data not shown), indicating that these cells may have elevated translational mechanisms. Lastly, the MuSC1 population has enriched expression of the myogenic gene *PAX7* and, to a lesser extent, *MYF5*, compared the MuSC2 population. These observations suggest that MuSC1 is comprised of quiescent MuSCs and MuSC2 is comprised of an early-activated MuSCs.

We performed Ingenuity Pathway Analysis (IPA) to compare biological processes differentially activated between the MuSC1 and MuSC2 populations. The IPA gene group “Oxidative Phosphorylation” is enriched in MuSC1 (Ryall et al., 2015), while “EIF2 Signaling”, associated with protein translation processes, is enriched in MuSC2 (**Fig. 2C**). Furthermore, Gene Set Enrichment Analysis (GSEA) also found MuSC1 to be enriched for “myogenesis”, “muscle cell differentiation”, “hypoxia”, and “response to mechanical stimulus” gene sets, consistent with the observation that these cells are both less differentiation and may have stress-associated gene induction due to tissue dissociation (van den Brink et al., 2017) (**Fig. 2D**). MuSC2 cells are enriched for “ribosome and translational initiation”, “MYC targets” and “E2F (cell proliferation)”, “G2M checkpoint (cell division)”, and “inflammation” gene sets, further supporting the interpretation that these cells may be in an early activated or partially differentiated state within an inflammatory environment (**Fig. 2D**). Taken together, these observations suggest that the MuSC1 population is comprised of quiescent MuSCs, while the MuSC2 population is comprised of active, proliferating, and/or dysregulated MuSCs, with expression alterations associated with inflammation, ageing, and muscle wasting. Differentially expressed genes such as *IL32*, *CXCL1*, *CCL2*, and *TNFRSF12/FN14* may constitute a marker set for MuSC variation in chronic muscle inflammation in various pathologies.

### Ligand-receptor interaction model identifies potential surface markers and cell-communication channels in muscle pathologies

Given the distinct expression profiles between the MuSC1 and MuSC2 populations, we sought to identify genes that could facilitate surface antigen-based separation of these two human MuSC populations for prospective isolation strategies. We identified surface receptor genes that were differentially expressed between the MuSC1 and MuSC2 populations, using a database of 542 human surface “receptor” genes (Ramilowski et al., 2015) (**Fig. 3A**). MuSC1 were exhibit elevated expression of *EGFR*, *ITGB1*, *FGFR4*, *SDC2*, as well as the three tetraspanins *CD81*, *CD82*, and *CD151*. EGFR is a recently established human MuSC marker and is required for basal-apical asymmetric cell division (Charville et al., 2015; Wang et al., 2019). The tetraspanin CD82 is also a recently recognized human MuSC maker (Alexander et al., 2016), while CD9 and CD81 have been identified to control muscle myoblast fusion (Charrin et al., 2013). Furthermore, Syndecans (SDCs) have been identified in mouse to be heterogeneously expressed on MuSCs and myoblasts during muscle repair (De Micheli et al., 2019) and have been shown to form co-receptor complexes with integrin β1 (ITGB1) and FGFR4 upstream of signaling pathways regulating myogenesis (Pawlikowski et al., 2017). Only SDC4 and SDC3 have yet been identified to mark adult mouse MuSCs (Pisconti et al., 2012). In comparison, the MuSC2 population has elevated expression of *CD44* and *TNFRSF12*/*FN14* as previously noted. The CD44 receptor has been shown to regulate myoblast migration and fusion in mouse, but also mark MuSCs in osteoarthritis patients (Mylona et al., 2006; Scimeca et al., 2015).

**Figure 3.**
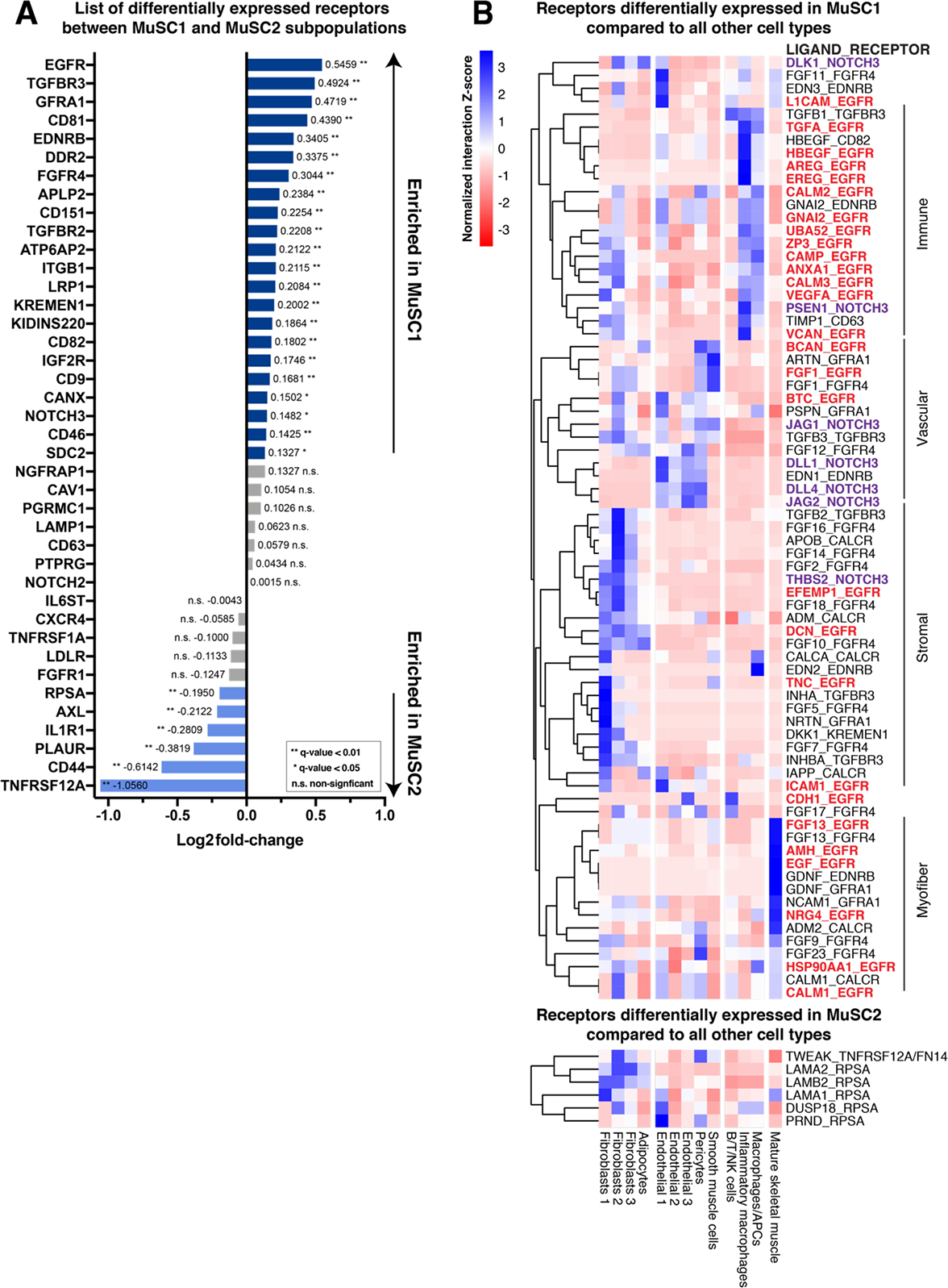
Differentially expressed receptors and ligand-receptor interaction scores. **(A)** List of differentially expressed genes between the MuSC1 and MuSC2 subpopulation ranked by log_2_ fold-change in expression. Positive average values correspond to genes that are upregulated in MuSC1, whereas negative values are upregulated in MuSC2. Receptors that are statistically significant (q-value is corrected for FDR < 0.05) are colored in blue. Receptors that are not statistically significant are in grey. **(B)** Heatmap representing row-normalized (Z-score) LR interaction scores. Rows represent ligand-receptor interaction pairs in the format LIGAND_RECEPTOR, where the receptor is either differentially expressed in the MuSC1 or MuSC2 populations compared to all the other cell types. Columns identify cell types expressing the ligand. Asterisks after the pair name also indicates that the ligand is differentially expressed by the other cell type and that interaction is likely cell-type specific. Red pairs involve the *EGFR* receptor, purple pairs the *NOTCH3* receptor. A positive value indicates that the interaction has a high score for a particular ligand and cell type compared to other cell types.

Next, we used a ligand-receptor (LR) interaction model and a database of LR pairs (Ramilowski et al., 2015) to map cell signaling communication channels in human muscle and uncover differences between the MuSC1 and MuSC2 populations. The model also identifies interacting ligand(s) and is restricted to receptor genes is differentially expressed by a specific cell type within the consensus human muscle cell atlas (**Fig. 1B**). For each LR pair, the model calculates an interaction score from differentially expressed receptors on MuSCs and ligands expressed by other cell types. We identified 73 and 6 significant LR interactions for the MuSC1 and MuSC2 populations, respectively (**Fig. 3**). Over one third of all interactions in the MuSC1 population involve the *EGFR* receptor, which has recently been shown to play a critical role in directing MuSC asymmetric division in regenerating muscle (Wang et al., 2019). A limited number of EGFR ligands have been identified in muscle repair. For example, amphiregulin (AREG) is secreted by T_reg_ cells (Burzyn et al., 2013). According to our model findings, EGFR may also interact with ligands expressed by immune cells, such as with TGF-α (*TGFA*), heparin-biding EGF (*HBEGF*), amphiregulin (*AREG*), and epiregulin (*EREG*). Other EGFR ligands include brevican (*BCAN*), and betacellulin (*BTC*) produced by endothelial cells, ECM proteins fibulin 3 (*EFEMP1*), decorin (*DCN*), and tenascin C (*TNC*) expressed by fibroblasts, and FGF13, AHM, NRG4 and EGF, expressed by mature skeletal myofibers. We also detect seven interactions involving *NOTCH3* with a variety of ligands. Notch3 signaling is involved in maintaining MuSC quiescence, in particular through interaction with DLL4 (Low et al. 2018), which we found differently expressed by endothelial cells along with *JAG2*. In addition, *NOTCH3* also interacts with the ECM protein thrombospondin-2 (*THBS2*).

Only two receptors, *TNFRSF12*/*FN14* and *RPSA*, were found differentially expressed in MuSC2 compared to other cell types. The first, *TNFRSF12*/*FN14*, interacts with the *TWEAK* cytokine ligand. While typically recognized to be expressed by macrophages and other immune cells (Tajrishi et al., 2014), our model suggests *TWEAK* is also expressed by the Fibroblasts 2 and Pericyte cell populations, though not in a statistically significant manner. The second, *RPSA*, is surface ribosomal protein that interacts with laminins (*LAM*), a dual-specificity phosphatase 18 (*DUSP18*), and prion protein 2 (*PRND*), which taken together may suggest various pathological processes such as prion diseases and cancer (Pampeno et al., 2014; Wu et al., 2019). Together, this ligand-receptor analysis identifies a broad set of surface markers that could refine the molecular definition of human MuSCs and their subpopulations, as well as candidate cell-communication channels differentially involved in healthy and diseased muscle tissues.

## Discussion

Here we present an annotated multi-donor dataset consisting of 22,000 single-cell transcriptomes from 10 different donors and unique anatomical sites, some of which difficult to access outside of reconstructive surgeries. We performed single-cell RNA sequencing and the bioinformatic integration method Scanorama to examine the cellular heterogeneity across diverse adult human muscle tissue samples. We observed that Scanorama performed data integration more successfully than other approaches (**Fig. S1**). We describe the muscle tissue cellular heterogeneity and provide a comprehensive analysis of differentially expressed genes for 16 resolved cell subpopulations (**Fig. 1**). This analysis suggests new gene markers for muscle FAPs and vascular endothelial cells that may provide unique perspective to human muscle physiology.

Most notably, this analysis suggests that human muscle may contain two distinct MuSC subpopulations (**Fig. 2**). Given the broad donor age range in this study, these two subpopulations may constitute a healthy and an aged/diseased MuSC pool. We conclude that the “MuSC1” subpopulation to be largely comprised of “quiescent” MuCSs, owning high levels of *PAX7*, the mitotic inhibitor *CDKN1C*, and *DLK1*. Interestingly, *DLK1* may be an important regulator for human MuSC maintenance and a marker of healthy tissue given its role in inhibiting adipogenesis (Andersen et al., 2013). Conversely, we identified in the “MuSC2” population signatures of inflammation and increased fat metabolism (*CCL2* and *CXCL1*), reduced insulin sensitivity (*IL32*), cell cycle (EIF2 Signaling terms), and muscle wasting (*TNFRSF12*/*FN14*), thereby suggesting that these cells may constitute an “early-activated” and possibly dysfunctional MuSC pool. These markers are consistent with prior observations that excessive fat accumulation in muscle can be attributed to obesity, diabetes, and ageing (Addison et al., 2014). In addition, we identify two upregulated lncRNAs that warrant further investigation as candidate non-coding regulators of myogenesis (Hagan et al., 2017). Moreover, the finding of two human MuSC subpopulations mirrors similar observations made from mouse muscle scRNA-seq analyses (De Micheli et al., 2019; Dell’Orso et al., 2019) and agrees with the general conceptual framework that muscle stem cells transition between quiescent, activated and cycling states (Bentzinger et al., 2013). Future studies comparative analysis of these MuSC subpopulations across species may reveal human-specific aspects of myogenesis. Ligand-receptor interaction models from scRNA-seq data can help formulate new hypotheses about cell-communication channels that regulate muscle function (De Micheli et al., 2019). Identifying new MuSCs surface receptors will also help us refine MuSC purification protocols for prospective isolation studies used for *in vitro* and transplantation models. Our LR model revealed a set of 40 surface receptor genes that are distinctly expressed between MuSC1 and MuSC2, confirming some prior reports but also providing new candidate surface antigens for human MuSC subpopulation fractionation (**Fig. 3**). For example, we identify that *SDC2* may mark “quiescent” MuSCs while *CD44*, *TNFRSF12*, and *RPSA* “early-activated” MuSCs in ageing and disease contexts. In addition, our model proposed 79 cell-communication signals that may act between MuSCs and other cell types; in particular with fibroblasts, myofibers and immune cells through the EGFR receptor, and with vascular cells through the NOTCH3 receptor. These interactions may be critical regulators of muscle homeostasis and should be further investigated.

Our study has some limitations. First, the sample size is small, and donors are very diverse, thus limiting our ability to control for age and sex. We performed differential expression and gene set enrichment analyses within the MuSC1 and MuSC2 populations between the middle-age (43-69 yo) and aged (70-81 you) donors, but found few age-cohort specific differences (data not shown). Nevertheless, our dataset still offers a new transcriptomic cell reference atlas and computational data integration approaches as a resource to examine human muscle cell diversity in health, ageing and disease.

Future studies should aim at collecting muscle specimens in a more controlled manner, for example using a Bergström needle (Tarnopolsky et al., 2011; Sarver et al., 2017) from a unique anatomical site; though this would not be possible for some muscles presented in this study. These biopsies would allow for ageing and disease comparative analyses. Indeed, a recent report by Rubenstein et al. (2020) performed scRNA-seq on four human vastus lateralis muscle biopsies found that myofiber type composition and gene expression alterations based on donor age. Further, future work could also focus on collecting single-myonuclei from myofibers while discarding other non-myogenic cell types. This could illuminate the transcriptomic diversity of myofiber type, on differences that the local anatomy and tissue physiology may demonstrate, and to ultimately enrich our repertoire of know human muscle markers and understanding of its molecular regulators.

## Methods

### Human participation for muscle sample collection

All procedures were approved by the Institutional Review Board at Weill Cornell Medical College (WCMC IRB Protocol # 1510016712) and were performed in accordance with relevant guidelines and regulations. All specimens were obtained at the New York-Presbyterian/Weill Cornell campus. All subjects provided written informed consent prior to participation. Samples were de-identified in accordance to IRB guidelines and only details concerning age, sex, and anatomic origin were included. Sample anatomic locations and donor details are provided in **Fig. 1A**.

### Muscle digestion and single-cell sequencing library preparation

After collection from donors during surgery, the muscle samples were cleared from excessive fat and connective tissue and weighted. About 50-65 mg of tissue was then digested into a single-cell suspension following a previously reported protocol (Spinazzola et al., 2017). Briefly, the specimen was digested in 8 mg/mL Collagenase D (Roche) and 4.8 U/mL Dispase II (Roche) for 1 hr followed by manual dissociation, filtration, and red blood cell lysis. All single-cell suspensions were then frozen at −80°C in 90% FBS, 10% DMSO and were re-filtered after thawing and prior to generating scRNA-seq libraries. The sequencing libraries were prepared using the Chromium Single Cell 3’ reagent V2 or V3 kit (10X Genomics) in accordance with the manufacturer’s protocol and diluted as to yield a recovery of ∼6,000 single-cell transcriptomes with <5% doublet rate. The libraries were sequenced in multiplex (n=2 per sequencing run) on the NextSeq 500 sequencer (Illumina) to produce between 200 and 250 million reads per library.

### Single-cell data analysis

Sequencing reads were processed with the Cell Ranger version 3.1 (10X Genomics) using the human reference transcriptome GRCh38. The downstream analysis was carried out with R 3.6.1 (2019-07-05). Quality control filtering, data clustering, visualization, and differential gene expression analysis was carried out using Seurat 3.1.0 R package. Each of the 10 datasets was first analyzed and annotated independently before integration with Scanorama (Hie et al., 2019). Filtering retained cells with >1000 unique molecular identifiers (UMIs), <20% UMIs mapped to mitochondrial genes, and genes expressed in at least 3 cells. Unsupervised shared nearest neighbor (SSN) clustering was performed with a resolution of 0.4 following which clusters were annotated with a common nomenclature of 12 cell type terms (**Fig. S1**). Differential expression analysis was achieved using either Seurat’s “FindAllMarkers” (**Fig. 1D**) or “FindMarkers” (**Fig. 2A**) function using a Wilcoxon Rank Sum test and only considering genes with >log2(0.25) fold-change and expressed in at least 25% of cells in the cluster. P-values were corrected for false-discovery (FDR) and then reported as q-values. Integration of raw counts was achieved using the “scanorama.correct” function from Scanorama. The integrated values were finally scaled in Seurat regressing out the 10X chemistry type and the number of genes per cell. Visualization was done using uniform manifold approximation and projection (UMAP) (Becht et al., 2018).

### Pathway and gene set enrichment analysis

The list of differentially expressed genes between MuSC1 and MuSC2 (Fig. 2A) was used in Ingenuity Pathway Analysis (IPA) (QUIAGEN, 2019-08-30). Activated (canonical) pathways were calculated by “Core Analysis” setting a q-value cutoff of 0.05, which yielded 964 genes (366 down, 598 up). Top canonical pathways were chosen based of -log(p-value) and z-score values. Gene set enrichment analysis (GSEA, v.4.0.3) (Subramanian et al., 2005) was ran on the same gene list as IPA ranked by log2 fold-change and with default program settings. Gene sets database used: h.all.v7.0.symbols.gmt, c2.all.v7.0.symbols.gmt, c5.all.v7.0.symbols.gmt (Broad Institute). Gene sets enriched in phenotype were selected based on q-value and enrichment score (ES).

### Ligand-receptor cell communication model

The model aims at scoring potential ligand-receptor interactions between MuSCs (receptor) and other cell types (ligand). We used the ligand-receptor interaction database from Ramilowski et al. (2015). From the database, we considered 1915 ligand-receptor pairs (from 542 receptors and 518 ligands) to test for differential expression in our scRNA-seq dataset. To calculate the score for a given ligand-receptor pair, we multiply the average receptor expression in MuSCs by the average ligand expression per other cell type. We only considered receptors that are differentially expressed in either the MuSC1 or MuSC2 subpopulation when compared individually to all other cell types.

## Reagents and Resources

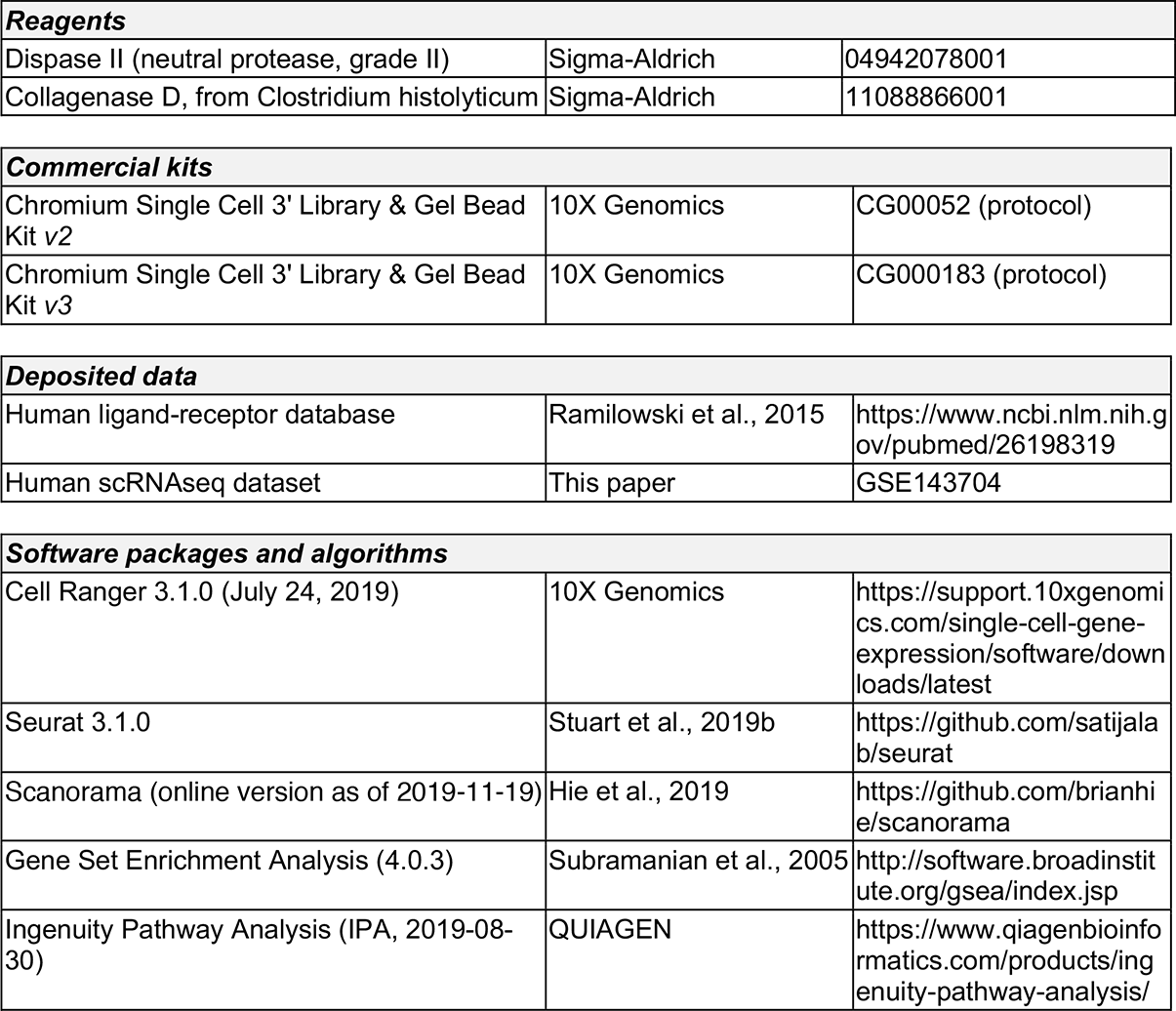

**Reagents**
|  |  |  |
| --- | --- | --- |
| Dispase II (neutral protease, grade II) | Sigma-Aldrich | 04942078001 |
| Collagenase D, from Clostridium histolyticum | Sigma-Aldrich | 11088866001 |
| **Commercial kits** |  |  |
| Chromium Single Cell 3' Library & Gel Bead Kit v2 | 10X Genomics | CG00052 (protocol) |
| Chromium Single Cell 3' Library & Gel Bead Kit v3 | 10X Genomics | CG000183 (protocol) |
| **Deposited data** |  |  |
| Human ligand-receptor database | Ramilowski et al., 2015 | https://www.ncbi.nlm.nih.gov/pubmed/26198319 |
| Human scRNAseq dataset | This paper | GSE143704 |
| **Software packages and algorithms** |  |  |
| Cell Ranger 3.1.0 (July 24, 2019) | 10X Genomics | https://support.10xgenomics.com/single-cell-gene-expression/software/downloads/latest |
| Seurat 3.1.0 | Stuart et al., 2019b | https://github.com/satijalab/seurat |
| Scanorama (online version as of 2019-11-19) | Hie et al., 2019 | https://github.com/brianhie/scanorama |
| Gene Set Enrichment Analysis (4.0.3) | Subramanian et al., 2005 | http://software.broadinstitute.org/gsea/index.jsp |
| Ingenuity Pathway Analysis (IPA, 2019-08-30) | QUIAGEN | https://www.qiagenbioinformatics.com/products/ingenuity-pathway-analysis/ |

## Acknowledgements

This work was financially supported by National Institutes of Health under award R01AG058630 (to B.D.C.), a Glenn Medical Research Foundation and American Federation for Aging Research Grant for Junior Faculty (to B.D.C.), and a US Department of Education Graduate Assistantship in Areas of National Need under Award P200A150273 (to A.J.D.). The content is solely the responsibility of the authors and does not necessarily represent the official views of any of these funding sources. The authors acknowledge helpful advice from colleagues in the Cosgrove and Elemento groups, as well as Christopher Mendias at the Hospital for Special Surgery and Peter Schweitzer of Genomics Facility at the Cornell University Biotechnology Resource Center. Lastly, the authors are grateful for the human tissue donors.

## Author Contributions

A.J.D. and B.D.C. designed the study and wrote the manuscript. J.A.S. obtained the human tissue samples. A.J.D. performed the tissue dissociations, scRNA-seq, and data analysis, with supervision and assistance from B.D.C. and O.E. All authors reviewed and edited the manuscript.

---

**Figure S1.**
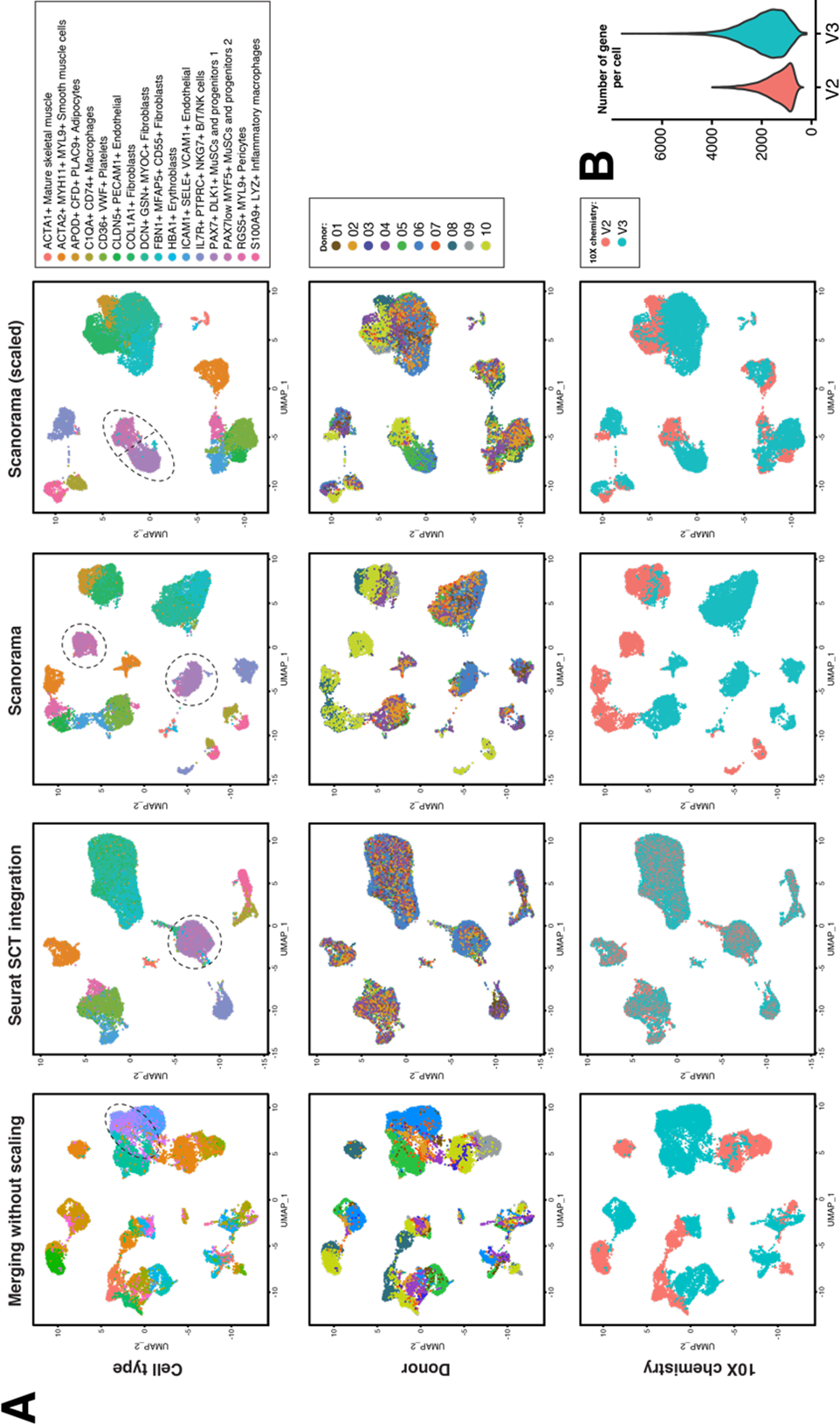
Comparison of scRNA-seq integration and batch correction methods. We compared four scRNA-seq data integration methods to evaluate which most faithfully conserves donor, anatomical, and biological information while minimizes technical biases. **(A)** The n=10 donor datasets were first annotated independently using a nomenclature of 12 common cell type terms following unsupervised SNN clustering. Then we evaluated the integration method by UMAP and by coloring the data either by cell type, donor ID, or 10X library chemistry used. *First*, we integrated the data by merging the individually normalized gene expression matrices without any further correction. We saw strong technical biases that overwhelmed biological information as the different cell populations segregate by sample/donor and chemistry type. For instance, the two MuSC and progenitor subpopulations are grouped with fibroblasts and endothelial cells. *Second*, we tested the Seurat SCT integration method (Stuart et al., 2019b). This method first calculates a cross-correlation subspace from genes that are shared between datasets. We noticed that this method better “aligns” donor and chemistry type but at the expense of masking biological variability. For instance, we observed that the two MuSC and four stromal subpopulations (Fibroblast 1,2,3 and Adipocytes) were grouped together, hiding important biological heterogeneity. Although certainly useful to validate reproducibility in scRNA-seq experiments, the Seurat SCT integration approach overcorrected biological heterogeneity for heterogeneous samples. *Third*, we tested the Scanorama method (Hie et al., 2019), which relies on a computer vision algorithm that “stitches” datasets together even when the cell type composition between dataset is considerably different. We see that this method groups similar cell populations together while acknowledging donor differences. Yet, surprisingly, this method is also very sensitive at picking up differences in chemistry. To correct this chemistry effect, we scaled the Scanorama output by regressing out the chemistry and the number of genes detected per cell (significantly different between chemistry type) **(B)**. Using this integration method, we observed clear separation of the independently annotated cell populations. We present the resulting Scanorama-integrated dataset as a “consensus atlas” (see **Fig. 1B-C**) of human muscle that describes donor-to-donor differences while grouping cells that are similar together and removing technical biases.

